# Unprecedented lead tolerance in an urban lizard

**DOI:** 10.1101/2024.12.06.626998

**Authors:** Annelise Blanchette, Kuan-Jiu Su, Alanna J Frick, Jordan Karubian, Alex R Gunderson

**Affiliations:** Tulane University, New Orleans, LA

**Keywords:** urban ecology, *Anolis*, heavy metals, locomotion, pollution, ecotoxicology, gene expression

## Abstract

Lead (Pb) is an extremely toxic heavy metal pollutant pervasive in many environments with serious health consequences for humans and wildlife. We found that Cuban brown anole lizards (*Anolis sagrei*) in New Orleans, USA, have the highest mean (955±877 µg/dL; N = 40) and individual (3,192 µg/dL) blood lead levels of any free-living vertebrate reported to date. Unexpectedly, this extreme field lead exposure did not decrease performance in whole-organism traits commonly affected by lead (balance, sprint speed, endurance). To identify toxicity thresholds, we employed a 60-day lead-dosing experiment and found that the brown anole blood lead threshold for decreased whole-organism performance is over an order of magnitude higher than the already extreme mean field concentration (10,600 µg/dL). Transcriptomic analysis of brain and liver from lizards in high and low lead sites revealed minor effects of lead exposure. However, several of the differentially expressed genes function in metal ion homeostasis and oxygen carrying capacity, pointing to cellular mechanisms that may contribute to high lead tolerance in these animals. The brown anole may be the most lead-tolerant vertebrate known to date and has the potential to serve as a powerful model system to help us understand mechanisms of lead tolerance.

## Introduction

Anthropogenic activity has resulted in extensive heavy metal contamination in water, soil, and air, contributing to significant declines in human and wildlife health (Ali and Khan 2019). Lead (Pb) specifically is a major concern due to its potent toxicity (Needleman 2004). Among many effects, lead is neurotoxic and genotoxic, causes neuromuscular disfunction and reduced cognition, diminished reproduction, anemia, and kidney, liver, and heart disease (Needleman 2004). Reflecting this toxicity, the California condor (*Gymnogyps californianus*) dwindled to near extinction due to lead poisoning in the 1980s (Finkelstein et al. 2012), and the Centers for Disease Control and Prevention (CDC) currently states that no level of blood lead is acceptable for children (CDC 2022). Lead use in paints, gasoline, and industrial processes has been greatly reduced in recent decades (Levin et al. 2021). However, lead does not break down and therefore many cities globally continue to have very high lead contamination due to historical deposition (Clune et al. 2011). As a result, finding organisms that can serve as biomonitors of lead contamination, as well as models of lead susceptibility and tolerance, remains a pressing issue (Tovar-Sánchez et al 2018).

We used the invasive Cuban brown anole lizard (*Anolis sagrei*) in New Orleans, Louisiana, USA, as a model system to investigate effects of urban lead contamination, integrating data on field lead exposure, experimental lead dosing, physiology, and functional genomics. New Orleans provides an ideal location of study because of decades of research into fine-scale geographic variation in soil lead contamination across the city (Figure 1A; Mielke et al. 2016). Soil lead in New Orleans covaries with tissue lead in humans and birds and is associated with detrimental outcomes (Mielke et al. 2016, McClelland et al. 2019, Hitt et al. 2023). Brown anoles are common in cities across the Southeastern United States, including New Orleans, are amenable to both field and laboratory study, and already serve as a model system in urban, physiological, and evolutionary ecology (Battles et al. 2019, Stroud et al. 2019, Deery et et al. 2021). Lizards are underrepresented in studies of heavy metal pollution, especially with respect to lead (Grillitsch and Schiesari 2016). Tissue-environment lead associations are known in some species (Burger et al. 2004, Nasri et al. 2016), but physiological studies of lead exposure in lizards are rare (Holem et al. 2006, Salice et al. 2009, Grillitsch and Schiesari 2016, Indest et al. 2018). Therefore, tissue lead toxicity thresholds in lizards, and by extension their toxicity thresholds relative to better-studied taxa such as birds and mammals, is poorly known.

**Figure 1.**
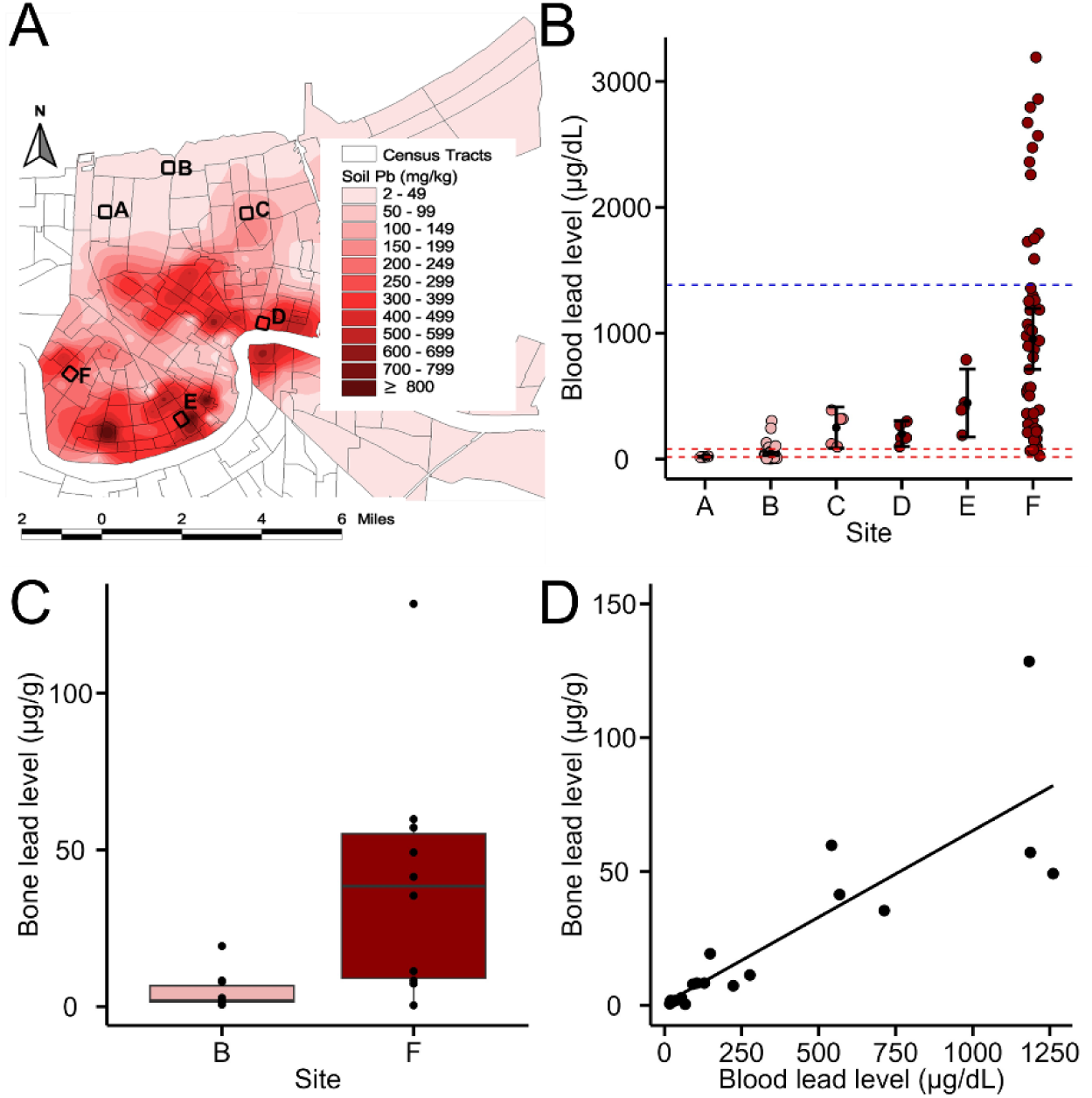
Soil and lizard lead levels in New Orleans. (A) Soil lead concentrations in New Orleans^8^ (Mielke et al. 2016) with our study sites indicated. (B) Blood lead concentrations of brown anoles by study site. Dashed red lines: reported whole-organism physiological toxicity thresholds for birds (71 µg/dL) and mammals (18 µg/dL; Buekers et al. 2009). Blue dashed line: the highest individual blood lead level from a live free ranging vertebrate that we could find in the literature (1384 µg/dL; griffon vulture (*Gyps fulvus;* Carneiro et al. 2016). (C) Bone lead levels of lizards collected from Sites B and F. (D) Relationship between bone and blood lead levels for the subset of individuals from Sites B and F for which we have both measures.

To test for an association between environmental lead contamination and lizard lead exposure, we measured blood lead concentrations of brown anoles from sites that vary in soil lead level across New Orleans. To determine if field lead exposure affects lizard whole-organism physiology, we tested for associations between blood lead level and multiple metrics of locomotor performance in field-caught animals. To experimentally determine blood lead toxicity thresholds, we measured multiple aspects of brown anole physiology during a 60-day lead dosing study in the laboratory. Finally, to investigate the effects of lead, and potential mechanisms of lead tolerance, at the molecular level, we sequenced transcriptomes of brown anoles from high and low lead contamination areas. We found that brown anoles in New Orleans have extreme blood lead levels and lead tolerance, while gene expression patterns highlight molecular mechanisms that may contribute to tolerance. Brown anoles may therefore be the most lead tolerant vertebrate known to date and provide a compelling vertebrate model system to investigate mechanisms of heavy metal tolerance.

## Methods

### Field Sampling and Animal Husbandry

Brown anoles were captured by hand in New Orleans, Louisiana and transported to Tulane University. Animal were housed individually in plastic cages (23 L x 15 W x 16 H cm) with a wooden dowel provided as a perch in fluctuating temperature (35/24°C, day/night) and humidity (60% daytime, 75% nighttime) controlled Percival chambers on a 14:10 light:dark cycle, conditions that mimic an average summer day in New Orleans (Deery et al. 2021). Lizards were watered twice daily and fed crickets dusted with calcium powder three times per week.

### Tissue Lead Analysis

We measured lead levels in the blood and bone of field animals. Blood samples (∼75 µl) were collected from the post-orbital sinus using heparinized capillary tubes (Kimble Chase; Sharma et al. 2014). Each sample was immediately transferred to a trace-metal-clean microcentrifuge tube and stored at −20°C until processing. For bone collection, animals were euthanized via cervical dislocation and whole femur, tibia, and fibula were dissected out and stored in a trace metal clean micro-centrifuge tubes at −20°C until processing.

Tissue lead levels of lizards collected from 2019 – 2021 were measured at the UC Santa Cruz Plasma Analytical Laboratory following established trace metal clean techniques and using ultra-pure reagents as outlined in Hitt et al. (2023). Blood samples were weighed and then dried overnight at 60°C to obtain dry weight, then cold digested for 10 hours in 100 µL (blood volume <50 µL) or 150 µL (blood volume >50 µL) of concentrated HNO_3_ (optima, Fisher Scientific). 30% H_2_O_2_ (ultrex, JT Baker) and ultrapure water was added to each sample for a HNO_3_ to H_2_O_2_ ratio of 2:1 and a final HNO_3_ concentration of 6%. Samples were vortexed and left to sit overnight before analysis. Lead concentrations were determined by inductively coupled plasma mass spectrometry (ICP-MS), measuring masses of ^208^Pb and ^205^Tl isotopes (used as an internal standard). For quality control, approximately 20 µL of NIST SRM 955c (lead in blood, level 2) was digested using the methods described above with an average recovery of 94% ± 3.4% RSD (N = 4). Frozen bone samples were thawed overnight then combined for each individual and weighed. They were digested for 6 h with sub-boiling concentrated HNO_3_ in closed Teflon vials then evaporated to dryness and diluted with 5% HNO_3_. Lead concentration was then analyzed as described above.

Blood lead levels in lizards collected in 2022 and 2023 were analyzed at the Louisiana Animal Disease Diagnostic Lab. 15 µL of blood were diluted in a solution of 0.1% Triton-X100 acidified to 0.2% with HNO_3_, resulting in a 21-fold dilution. Seronorm-Trace Elements (lead in blood, level 2, 303 ng/ml Pb) was used as a standard reference material and digested using the methods above with an average recovery of 103% RSD. Clotted samples were first weighed and digested with 600 µL of 35% Trace Metal HNO_3_ overnight at 85°C. Samples were then diluted to a 1/60 solution with 0.2% HNO_3_. Lead concentrations were determined using a Perkin Elmer PinAAcle 900Z Zeeman graphite furnace spectrometer.

### Whole Organism Performance

With field-collected animals, we tested for associations between individual blood lead level and performance in whole-organism traits known to be affected by lead exposure in other vertebrates: balance, sprint speed, and endurance (Yu et al. 1996, Burger et al. 2004, Mansouri and Cauli 2009, Mansouri et al. 2013). To measure balance, lizards were run on a 1.2 m long horizontal wooden dowel (Spezzano and Jayne 2004) covered with fine mesh. We counted the number of times an individual slipped, defined as anytime the hind legs lost traction such that the pelvis moved to the side of the dowel (Spezzano and Jayne 2004). Runs were filmed using an Apple iPhone 8 Plus. Lizards were run twice with 15 min between runs. The dowel was 1.2 m off the ground and foam was placed underneath in case individuals fell or jumped. The diameter of the dowel was 0.79 cm or 0.64 cm for lizards with a knee-to-knee width of ≥ 27 mm and ≤ 26 mm, respectively, to account for inter-knee distance (Spezzano and Jayne 2004). Trials were conducted at 32°C, the body temperature of maximum field activity for *A. sagrei* in New Orleans (Ryan and Gunderson 2021).

Sprint speed was measured by video analysis of lizards running up a 1.8 m wooden track set at a 37° angle and covered with fine mesh for traction (Gunderson et al. 2018). Perpendicular lines were drawn every 12.5 cm and the fastest interval was taken as an individual’s maximum speed for each run. The average of the two fastest intervals was used in analyses. Each lizard was run twice at 32°C and filmed at high speed (120 frames per second) with an Apple iPhone 8 Plus. Runs were 15 min apart.

We measured endurance as time to exhaustion (s) while lizards ran at 32°C on a motorized treadmill (DogPacer MiniPacer) at 0.80 km^−1^ (Lailvaux et al. 2018). Lizards were placed on the treadmill while tethered with thread around the waist and induced to run by gently tapping their tails. Exhaustion was indicated by a loss of righting response. Endurance was measured twice per lizard with 24h between trials and the mean endurance time was used in further analysis.

### Lead Dosing Study

In 2023, N = 50 males were captured at low lead contamination Site B (Figure 1). After one week of lab acclimation, individuals were randomly assigned to one of five treatment groups (N = 10/treatment). Treatments were administered daily by oral gavage (Holem et al. 2006, Salice et al. 2009) for 60 days. The control group received distilled water, while the treatment groups received 1, 10, 100, or 500 mg lead/kg bodyweight, respectively. Following methods previously used in lizards (Holem et al. 2006, Salice et al. 2009, Indest et al. 2018), lead solutions were prepared by diluting lead acetate trihydrate (ThermoFisher Scientific Chemicals) in distilled water (ThermoFisher Scientific Chemicals). We measured seven traits/endpoints: symptomatic lead poisoning, endurance, sprint speed, balance, kidney size, liver size, and testis size (Yu et al. 1996, Burger et al. 2004, El-Said et al. 2008, Mansouri and Cauli 2009, Salice et al. 2009, Gonick 2011, Mansouri et al. 2013, Hora and Wuestefeld 2023). Symptomatic lead poisoning was defined as loss of appetite for one week and/or presence of oral ulcers (El-Said et al. 2008). Animals were removed from dosing if they showed symptomatic lead poisoning. Endurance was measured three times per lizard using the methods described above: the day prior to dosing, after 30 days of dosing, and after 60 days dosing (i.e., the end of the experiment). Individuals removed early due to lead poisoning had their endurance measured on the day of removal. Sprint speed and balance were measured as described above at 60 days on individuals in the control, 1 mg/kg, and 10 mg/kg treatment groups. Blood samples were collected from all individuals prior to the beginning of dosing and after all performance measurements at the end of dosing. Blood lead level did not differ among groups prior to dosing (F-value = 1.05, P = 0.394). Animals were anaesthetized with isoflurane and euthanized via cervical dislocation, after which the liver, kidney, and testes were removed and wet weight was measured with a digital balance.

With our dosing data, we calculated the blood lead toxicity threshold for a trait as the highest blood lead level of an individual within the No Observed Effect Concentration (NOEC) treatment for that trait. NOEC is defined as the highest lead dose that does not cause a significant change from the control group (Buekers et al. 2009).

### Statistical Analyses of Lead and Performance Data

All statistical analyses were performed using R Statistical Software (v4.1.2; R Core Team 2021). For the field collected animals, we tested for differences in blood and bone lead level between lizards from high and low lead sites using multiple linear regression with site soil lead level (‘high’ or ‘low’) as a predictor variable, with sex and site by sex interaction covariates for blood analyses and sex and snout-vent-length (SVL) as covariates for bone analyses. We tested for an association between blood and bone lead level within individuals with a Pearson product-moment correlation test. We tested for associations between individual blood lead level and endurance and sprint speed with multiple linear regression with sex and SVL as covariates. For tests with balance, negative binomial models were used with sex and relative leg length (residual knee-to-knee width relative to SVL) as covariates.

For the dosing study, we first tested for differences in blood lead level among treatment groups prior to dosing with a one-way ANOVA (there were none; P = 0.394). We tested for differences in blood lead level within and among treatment groups due to dosing using a two-way repeated measures ANOVA. One individual in the 500 mg/kg treatment group was found deceased on day 10 of dosing and was excluded from the analyses since a second blood sample could not be collected. We tested for effects of lead dosing on physiological performance at the treatment and individual level. For treatment-level analyses, treatment group was a predictor variable, and for individual-level analyses, individual blood lead level was a predictor variable. For endurance, linear mixed effects models were used with endurance as the response variable, day of measurement and SVL as covariates and individual as a random effect. Two individual-level analyses were conducted for endurance, one with data from all treatments including animals that were removed early due to lead poisoning, and another with only the animals that made it through the entire study. We also tested for differences in sprint speed and balance at the end of the study among the Control, 1 mg/kg, and 10 mg/kg animals at the treatment and individual levels. We tested for differences in balance among treatment groups using a general linear model with a Poisson distribution. We tested for differences in liver, kidney, and testis mass among treatments using one-way ANOVAS. Organ masses were normalized to body mass (organ wet mass/total body mass) before analysis.

### Transcriptome Analysis

To investigate the cellular processes affected by lead exposure, and to provide insight into how the lizards may tolerate high lead levels (see Results), we tested for differential gene expression of lizards collected from a high (Site F) and a low (Site B) lead contamination area in 2021 (Figure 1; N = 5 males and N = 5 females per site). Forebrain (males) and liver (males and females) transcriptomes were sequenced. We chose brain due to the neurotoxicity of lead and liver due to its role in detoxification (Needleman 2004, Hora and Weustefeld 2023). Forebrain was only collected from males due to small female size. After one week in the laboratory under our standard housing regime (see above), individuals were anaesthetized as described above and the target tissues were immediately dissected out, placed in RNAlater, flash frozen in liquid nitrogen, and stored at −80°C. Total RNA was extracted using the Qiagen RNeasy Plus Universal Tissue Mini Kit. Transcriptome libraries were prepared and sequenced at Novogene Corporation Inc (Sacramento, CA) using 150-base pair paired-end sequencing on the Illumina NovaSeq 6000 platform. Sequencing resulted in an average of 25,569,255 ± 3,909,485 reads per transcriptome.

We used Rcorrector to correct random sequencing errors in reads and removed kmers with errors that were unfixable (Song and Florea 2015). Trimmomatic was used to filter the resultant sequences based on quality, which was assessed for each read with a 4bp sliding window (Bolger et al. 2014). Sequences were trimmed when Phred33 quality scores fell below an average of 15 within the window. Adaptor sequences, if detected, were removed. Quality-controlled sequence reads were mapped to the *Anolis sagrei* genome (AnoSag2.0; Geneva et al. 2021) using STAR (Dobin et al. 2013).

We used the DESeq2 package (Love et al. 2014) to identify differentially expressed genes (DEG) between high and low lead groups based on sex and tissue type. We filtered out low expression genes, and only conducted analyses on those with a count threshold of 10 in at least three samples (Love et al. 2014). False discovery rate (FDR) correction for multiple comparisons was applied to maintain a table-wide alpha of 0.05 (Storey and Tibshirani 2003). Genes were considered differentially expressed when the absolute value of log2 fold expression difference between high and low lead groups was ≥ 1 and FDR was <= 0.05 (Love et al. 2014). Volcano plots were generated using EnhancedVolcano (version 1.16.0; Blighe et al. 2022). Gene Ontology enrichment analyses were conducted on differentially expressed genes using Enrichr (Kuleshov et al. 2016).

## Results and Discussion

### Field Lead Exposure

Lizard blood lead levels varied with soil lead contamination across New Orleans (Figures 1A, B). In the two sites with the most data (low lead contamination Site B and high lead contamination Site F; N = 40/site), blood lead in the high contamination site (x̅ ± SD; 955±877 µg/dL) was over 23 times higher than that of lizards from the low contamination site (40.4 ± 50.6 µg/dL; P = < 0.001; Figure 1B; Table S1). Additionally, blood lead levels across the city frequently exceeded the reported toxicity thresholds (Buekers et al. 2009) for whole-organism physiological effects in birds (71 µg/dL) and mammals (18 µg/dL; Figure 1B). The highest individual blood lead level was 3192 µg/dL, which is 2.3 X higher than the highest individual blood lead levels that we could find reported for an individual free-ranging vertebrate, a griffon vulture (*Gyps fulvus*) that died within 24 hours of capture (Carneiro et al. 2016; Figure 1B).

Blood lead reflects shorter-term exposure because lead is absorbed into the bloodstream within hours of ingestion and has a half-life that ranges between 3 and 30 days in wildlife and humans (Elder et al. 2014). To determine if the lizards are exposed to lead over longer timescales we measured lead in bone, which begins incorporation during development and can remain for years without further exposure or continue to increase as exposure continues and further incorporation occurs (Needleman 2004). Bone lead was significantly higher in high contamination Site F (39.9 ± 38.1 µg/g) than in low contamination Site B (4.67 ± 5.83 µg/g; P = 0.012; Figure 1C), indicative of long-term exposure in high contamination areas. Furthermore, individual bone and blood lead levels were significantly correlated (*r* = 0.86, t_(18)_ = 7.16, P < 0.001; Figure 1D), consistent with lizards residing within their respective sites throughout their lives. Indeed, wild brown anoles rarely move more than a few meters from their natal site (Calsbeek 2009) and adults have small home ranges (mean = 17.6 m^2^ for males; Schoener and Schoener 1982). Such high philopatry and adult site fidelity is the norm for *Anolis* lizards generally (Schoener and Schoener 1982), and therefore these lizards (and perhaps lizards more broadly; Talent et al. 2002) are strong candidates to serve as biomonitors of environmental lead contamination.

To place the blood lead levels of brown anoles in New Orleans into a broader context, we compiled all mean blood lead levels we could find reported for live, free-living adult vertebrates (minimum sample size N = 10). If multiple values were found for the same species, only the highest mean value was included. Brown anoles in New Orleans are clearly exceptional (Figure 2). The mean lead level in Site F is over three times higher than the next highest mean value (Nile crocodiles, x̅ = 250 µg/dL; Humphries et al. 2022), and 10 times higher than the peak mean level in California condors that experienced population declines due to lead exposure (130 µg/dL; Finkelstein et al. 2012; Figure 2). How these lizards attain lead is currently unknown. High lead levels in crocodiles and scavenging birds like condors and vultures are linked to the ingestion of large pieces of lead associated with fishing and hunting equipment (Pain and Mateo 2019, Humphries et al. 2022). That pathway of lead exposure is not possible for brown anoles as small (adult mass = 1.5 – 8 g) insectivores that feed on live prey. Instead, they must ingest it from a combination of diet, drinking water, and breathing or ingesting lead-contaminated dust. Arthropods do accumulate lead in lead-contaminated areas (Khan 2019), although a significant portion of that lead may be stored in the exoskeleton in a form not bioavailable to predators (Inouye et al. 2007). Ingestion of lead in dust is a very common means of lead exposure for humans and wildlife (Mielke et al. 2016), and brown anoles spend the majority of their time near and on the ground in close proximity to contaminated soils.

**Figure 2.**
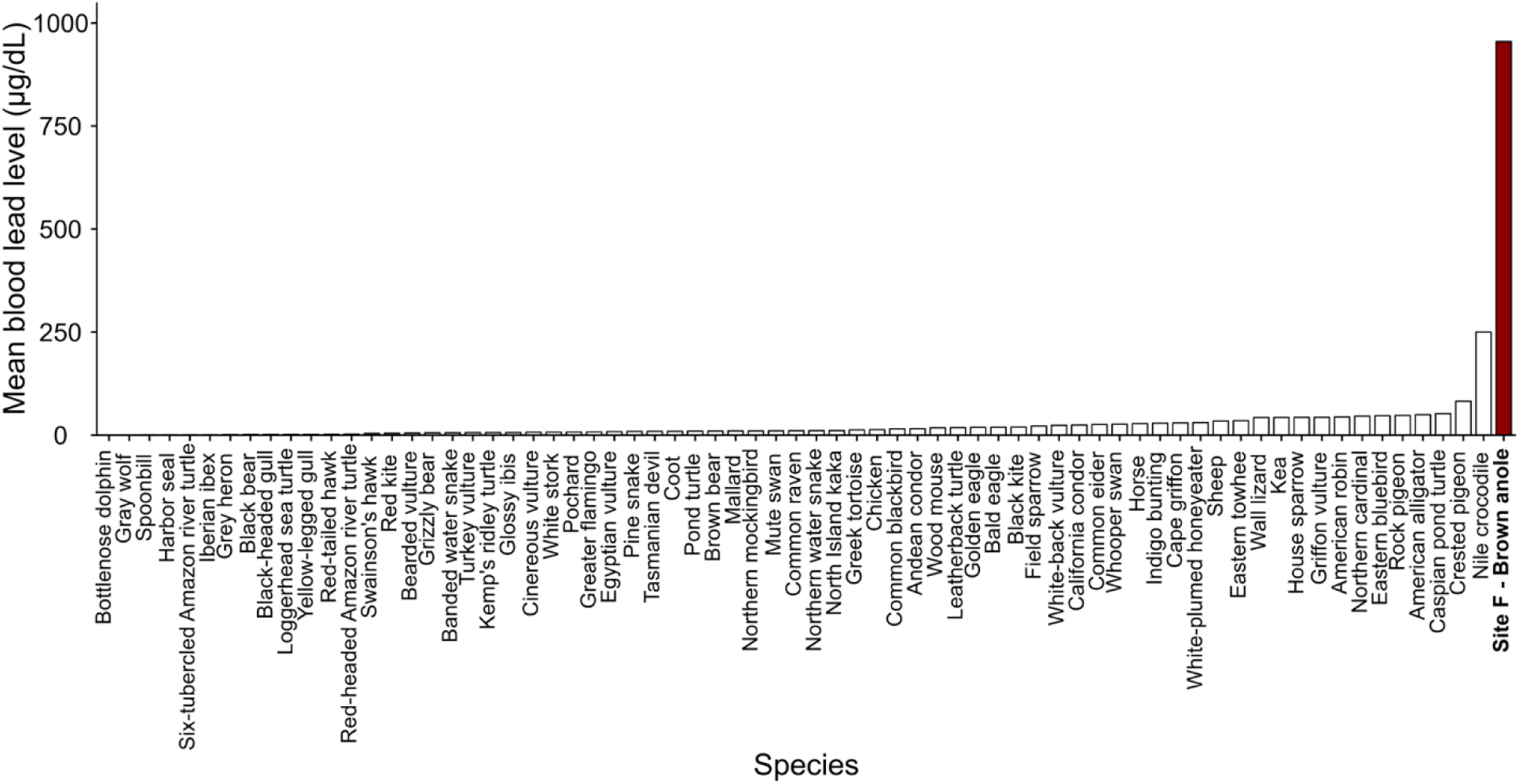
Mean blood lead concentrations of free-living vertebrates. The brown anole value is from high lead contamination Site F (Figure 1). Data for other taxa were reported in the literature (minimum N = 10). If multiple values for a species were found we included only the highest. See Dataset S1 and References S1 for more information.

### Physiological Performance of Field Exposed Lizards

Given their high lead levels, we predicted that field-exposed lizards would experience physiological detriments. Contrary to our expectation, blood lead was not associated with decreased performance (Figure 3). Blood lead was unrelated to balance (P = 0.276; Figure 3A; Table S2), positively related to sprint speed (P = 0.01; Figure 3B; Table S3), and trended towards a positive relationship with endurance (β = 0.04; P = 0.086 Figure 3C; Table S4). Brown anoles in New Orleans therefore appear to have greater lead tolerance than birds and mammals (Buekers et al. 2009). However, these data do not allow us to estimate their blood lead thresholds because no performance detriments were observed. To estimate brown anole lead toxicity thresholds, we turn to our experimental lead dosing experiment.

**Figure 3.**
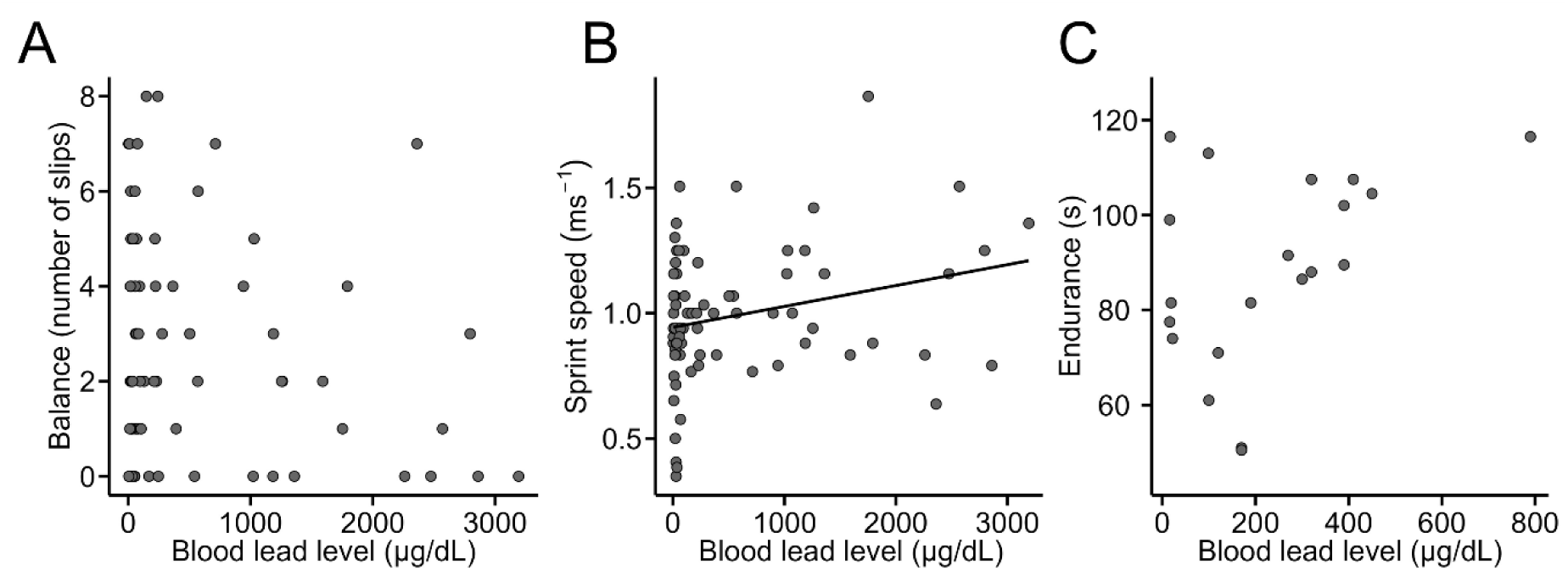
Physiological performance of individual field-caught brown anoles relative to blood lead level. (A) Balance. (B) Sprint speed. (C) Endurance.

### Effects of Experimental Lead Dosing

Blood lead level increased in all groups over time consistent with treatment level (Figure 4A, Tables S5, S6). The final mean blood lead level of lizards in all of the lead-dosed treatments was higher than the mean blood level at our most contaminated field site (Figure 4A, Table S5). All lizards in the 100 and 500 mg/kg treatments exhibited symptomatic lead poisoning before the end of the experiment and were removed from dosing by day 40 and day 10, respectively. In contrast, no lizards exhibited symptomatic lead poisoning in the Control, 1 mg/kg, or 10 mg/kg treatments. The NOEC for symptomatic lead poisoning is therefore 10 mg/kg, and the blood lead toxicity threshold is 10,600 µg/dL (Table 1).

**Figure 4.**
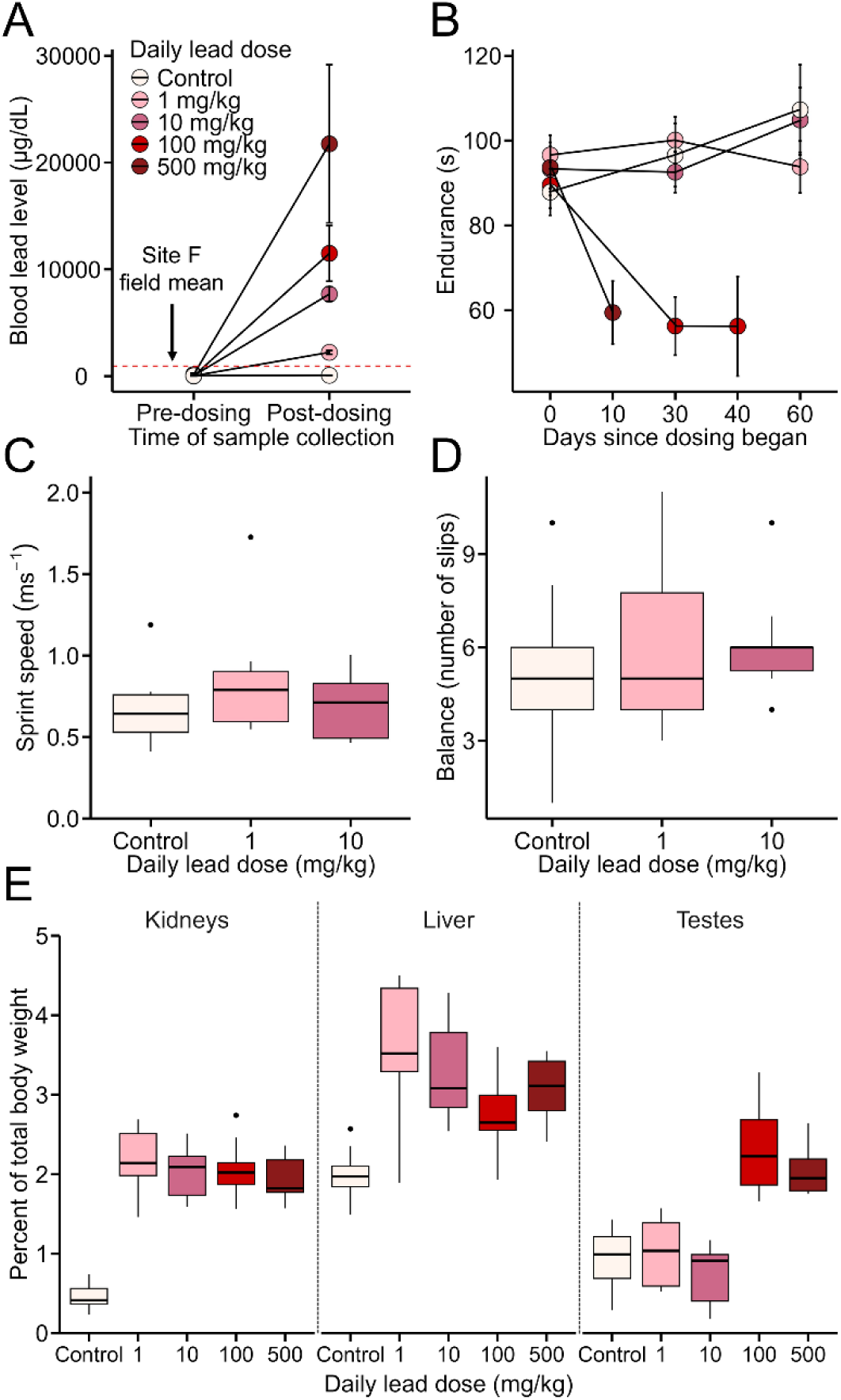
Effects of lead dosing on brown anole lizards. (A) Blood lead concentrations before and at the conclusion of dosing. Post-dosing samples were collected at 60 days except for the individuals removed early due to symptomatic lead poisoning in the 100 mg/kg and 500 mg/kg treatment groups. Points and bars indicate means and standard errors, respectively. The dashed line indicates the mean blood lead level of field-caught brown anoles from Site F, which had the highest lead level of all field sites. (B) Locomotor endurance by treatment group over time. (C) Sprint speed and (D) balance of lizards in the Control, 1 mg/kg, and 10 mg/kg treatment groups at the conclusion of dosing. (E) Organ wet weight as percent of total body mass by treatment.

**Table 1.** Brown anole lead toxicity thresholds from the dosing experiment. NOEC = No Observed Effect Concentration. Blood lead toxicity threshold = highest individual blood lead concentration in the NOEC treatment. We do not have a direct estimate of NOEC for kidney and liver size because both organs were enlarged in the lowest-dose treatment, and we report the highest individual blood lead concentration in the Control treatment as the toxicity thresholds for both.

| Endpoint | NOEC (mg/kg) | Blood lead toxicity threshold (µg/dL) |
| --- | --- | --- |
| Symptomatic lead poisoning | 10 | 10,600 |
| Endurance | 10 | 10,600 |
| Sprint speed | 10 | 10,600 |
| Balance | 10 | 10,600 |
| Kidney size | < 1 | 130 |
| Liver size | < 1 | 130 |
| Testis size | 10 | 10,600 |

Endurance differed by treatment (F = 4.81, P < 0.001), decreasing over time in the 500 mg/kg and 100 mg/kg treatments, but not in the others (Figure 4B). The NOEC for endurance is 10 mg/kg, and the estimated blood lead toxicity threshold is 10,600 µg/dL (Table 1). We also tested for an association between final individual blood lead level and final endurance. With data from all treatments, blood lead is negatively associated with endurance (blood lead range = 53 – 80,000 µg/dL; R^2^ = 0.47, β = −0.007, P < 0.001; Figure S1A, Table S7). However, when animals from the treatments above NOEC were excluded, endurance and blood lead were unrelated (blood lead range = 53 – 10,600 µg/dL; R^2^ = 0.05, β = 0.001, P = 0.432; Figure S1B).

Sprint speed and balance were unrelated to lead for the only animals that remained at the end of the experiment (the Control, 1 mg/kg, and 10 mg/kg groups) at both the treatment (both P > 0.325; Figures 4C, D; Tables S8, S9) and individual blood lead levels (both P > 0.781; Figures S1C, D; Tables S8, S9). We therefore cannot directly calculate NOEC or blood lead toxicity thresholds for these traits, though we can say that the NOEC and blood lead threshold are at least as high as for endurance (Table 1).

Given the dosing results, it is unsurprising that we found no whole-organism physiological detriments of what at first appeared to be very high levels of field lead exposure (Figure 3). The whole organism performance of these animals is not affected until blood lead concentration is much higher than field levels. In our site with the highest lead levels, Site F, the mean blood lead level was 955 µg/dL and the highest documented individual blood lead level was 3192 µg/dL (Figure 1). However, we only began to see whole organism performance effects for individuals in the 100 mg/kg treatment, which had a mean blood lead level 12x higher than the field mean (Figure 2A, Table S5).

All lead-dosed treatments had larger kidneys (F-value = 56.47, P < 0.001) and livers (F-value = 12.79, P < 0.001) than Controls, including the 1 mg/kg and 10 mg/kg groups that exhibited no whole-organism evidence of lead effects (Figure 4E). In contrast, testes were only enlarged in the 100 mg/kg and 500 mg/kg treatments (F-value = 29.50, P < 0.001; Figure 4E). The NOEC for effects on liver and kidney size are therefore lower than for whole-organism traits and cannot be directly estimated, but we know the NOEC must be less the 1 mg/kg (Table 1). We report the blood lead toxicity threshold as 130 µg/dL, the highest individual concentration in the Control group (Table 1). Whether or not the liver and kidney responses reflect an adaptive response to lead or pathology are unclear and require further investigation. The kidney and liver remove and sequester heavy metals (Gonick 2011, Hora and Wuestefeld 2023) and increased organ mass may improve detoxification (Hora and Wuestefeld 2023). However, enlarged organs can also reflect heavy metal-induced tissue damage (Habeebu et al. 2000, Gonick 2011). The response of testis mass to dosing was similar to that of whole-organism traits, and had the same thresholds (NOEC = 10 mg/kg; blood lead = 10,600 µg/dL; Table 1). Testes do not generally function in heavy metal detoxification or adaptive sequestration, so changes in testis mass likely reflects harm from lead exposure (Salice et al. 2009, Heidaeri et al. 2021).

### Differential Gene Expression of Field Exposed Lizards

The high lead tolerance of brown anoles raises the questions of how this tolerance is achieved. Lead exposure in vertebrates typically causes the differential expression of hundreds if not thousands of genes within a given tissue (Schneider et al. 2012, Shi et al. 2020). Genes commonly affected by heavy metal exposure are those related to the cellular stress response (e.g., antioxidants and molecular chaperones; Needleman 2004, Gupta et al. 2010), transport of key ions that heavy metals can replace within biological processes (Needleman 2004), and genes that function in the binding and sequestration of metals (Gonick 2011). Indeed, tolerance for lead in vertebrates has been linked to metallothioneins (MT), a family of metal-binding proteins that can function in sequestering lead in a nontoxic form (Gonick 2011).

We found relatively few genes differentially expressed (DE) in high versus low lead brown anoles. In male forebrain, there were only four DE genes in high lead lizards, two upregulated and two downregulated (Figure 5A). In male liver, 51 genes were DE, with 41 upregulated and 10 downregulated (Figure 5B). In female liver, 23 genes were DE, 14 upregulated and 9 downregulated (Figure 5C).

**Figure 5.**
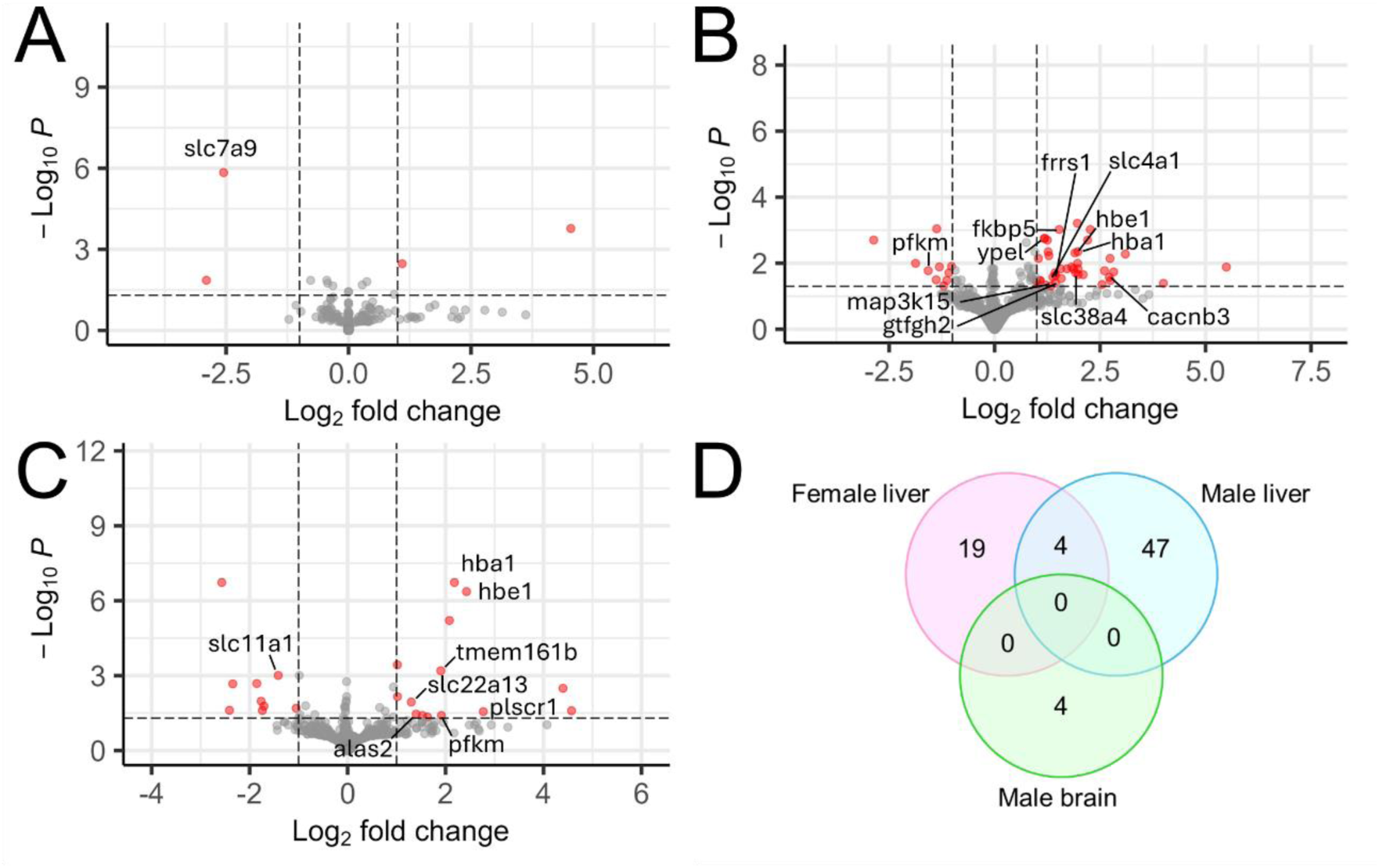
Differential gene expression of lizards from high compared to low lead contamination sites. Lizards were from high contamination Site F and low lead contamination Site B. Highlighted genes function in ion binding and transport, the oxidative stress response, or oxygen binding and transport. (A-C) Volcano plots of differential expression in the (A) male forebrain, (B) male liver, and (C) female liver. Cutoff for significance (red dots) is False Discovery Rate < 0.05 and an absolute log2 fold change ≥ 1. (D) Venn diagram of unique and shared differentially expressed genes across sex and tissue type.

There were no significantly enriched Gene Ontology (GO) categories across all DE genes after correction for multiple comparisons (Table S10). However, six Biological Processes neared significance (0.10 > P > 0.05), including Intracellular Iron Ion Homeostasis (GO:0006879) and Oxygen Transport (GO:0015671). Many of the DE genes are involved in the binding, transport, and homeostasis of metals and other ions (*alas2, frrs1, ypel3, plscr1, tmem161b, cacnb3, slc7a9, slc4a1, slc38a4, slc22a13, slc11a1*), and are also DE under heavy metal exposure in other taxa (He et al. 2009, Plusquin et al. 2012, Schneider et al. 2012, Truong et al. 2018, Smith et al. 2023). The response of several solute carrier genes (*slc*) is intriguing. SLCs are transmembrane proteins that function in ion transport into and out of cells and have been implicated in heavy metal susceptibility and tolerance (He et al. 2009, Schneider et al. 2012). We found no DE of metal-binding metallothionein genes.

One of the key consequences of lead exposure is anemia (Fuller and Wiley 2018) because lead interferes with heme production and decreases hematocrit, hemoglobin, and erythrocyte levels in vertebrates (Kwong et al. 2004), including western fence lizards (Salice et al. 2009). Key genes related to oxygen carrying capacity were upregulated in high lead anoles: *alas2* in female liver, and *hba1* and *hbe1* in both male and female liver (two of only four genes differentially expressed in both sexes). ALAS2 is a rate-limiting enzyme in heme production within red blood cells (Tanimura et al. 2016). *Hba1* produces alpha globin 1, one of two globins that form the mature hemoglobins that carry oxygen within vertebrate red blood cells. *Hbe1* is not part of primary hemoglobin function, but *hbe1* upregulation in mice ameliorates sickle cell disease and pathological underproduction of hemoglobin (thalassemia; Wang et al. 2016). Furthermore, there is evidence for parallel natural selection on the *hbe1* gene in populations of humans, dogs, yaks, and horses adapted to high-altitude hypoxia (Liu et al. 2019).

None of the top GO terms were associated with stress, and few stress-response genes were upregulated in high lead lizards. There were some key exceptions. *fkbp5*, *map3k15*, and *gtf2h2* are stress-related genes upregulated in male liver. *Fkbp5* produces an HSP90 cochaperone integral to the stress response (Gupta et al. 2010) and is also upregulated in carp cells in response to cadmium (Luan et al. 2022). *map3k15* is involved in stress-induced apoptosis, while *gtfh2* is involved in DNA repair (Tulay et al. 2015, Liu et al. 2022). *pfkm*, upregulated in female liver, can be induced by oxidative stress (Tang et al. 2012). None of the antioxidant genes frequently associated with heavy metal exposure, such as sodium oxide dismutase, catalase, or glutathione peroxidase, were DE.

Overall, we found minimal gene regulatory response to high levels of environmental lead exposure in brown anoles. Given the lack of higher-level physiological effects (Figure 3) this is perhaps not surprising, and further reinforces that these animals are very lead tolerant. Among the few genes that were differentially expressed, there are intriguing clues as to how the lizards tolerate lead. In particular, plasticity in the expression of solute transport genes and genes involved in oxygen carrying capacity are clearly candidates for further study.

## Conclusion

We found that brown anoles are reliable lead biomonitors (Burger et al. 2004) and provide evidence that they have extreme lead tolerance. Indeed, they may be the most lead tolerant vertebrate known. Many questions remain. Because of the scant attention paid to lizard heavy metal tolerance, we do not know if our findings reflect only brown anoles in New Orleans, brown anoles in general, the genus *Anolis*, or squamates more broadly (Moore et al. 2025). The high lead tolerance we observed could result from evolution in response to urban lead exposure. Heavy metal tolerance can evolve rapidly (Merritt and Bewick 2017), and *Anolis* lizards are known to quickly physiologically evolve in response to environmental change (Gunderson and Leal 2012, Campbell-Staton et al. 2017). Regardless, brown anoles represent a compelling, novel vertebrate model for developing a mechanistic understanding heavy metal tolerance with extensive resources available (Rasys et al. 2019, Geneva et al. 2021). Our findings also point to a need for greater consideration of heavy metals in the rapidly growing field of urban ecology and evolution (Rivkin et al. 2019). Cities across the globe have high levels of heavy metal pollution (Clune et al. 2011), but far more effort is devoted to factors such as urban heat islands, light and noise pollution, structural habitat modification, and invasive species (Rivkin et al. 2019). Taking steps to better understand the effects of heavy metals in urban areas, and how they interact with other well-known urban stressors, outside of a traditional ecotoxicological context will likely yield important insights (Schneeweiss et al. 2023, Sigmund et al. 2023).

## Supporting information

All Supplementary Materials

Supplemental Data for Figure 2

## Acknowledgements

We would like to thank R. Brasso, H. Frank and S. Chaturvedhi for guidance with this work, members of the Gunderson lab for their insights and assistance with field work, and M. Finkelstein and P. Jowett for assistance with tissue lead analyses. Funding was provided by The Morris Animal Foundation, the ASIH Gaige Fund Award, Sigma-Xi Grants in Aid of Research, and the Department of Ecology and Evolutionary Biology at Tulane. All work was approved by the Tulane University Institutional Animal Care and Use Committee (protocol #1436).

## Author Contributions

A.B., A.G. and J.K. conceived the project. A.B. and A.J.F. conducted physiological experiments. B. and K.S. conducted bioinformatic analyses. A.B. and A.G. wrote the paper with input from other authors.

## Data Availability

Supplementary information is available for this paper. Data presented here are available in a GitHub repository (https://github.com/AnolisNOLA/Brown-anole-data)

