## Supplementary material for "Unprecedented lead tolerance in an urban lizard": All Supplementary Materials

This PDF file includes:

Figure S1

Tables S1 to S10

References S1

Other supporting material for this manuscript includes the following:

Dataset S1

**Results**

**Figure**

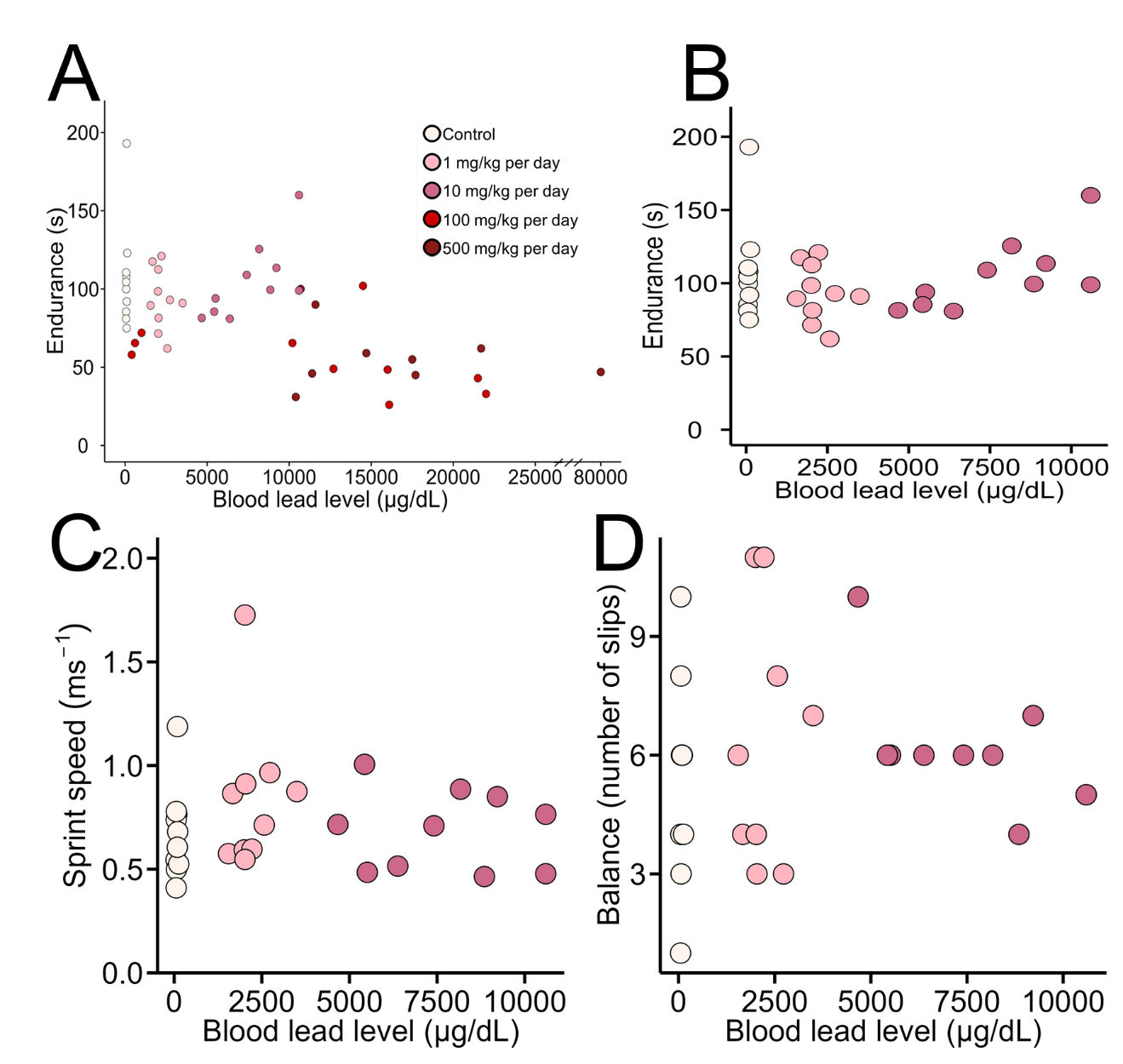

**Figure S1.** Physiological performance of lizards in the dosing study by blood lead level. (A) Final individual blood lead level was negatively associated with endurance when lizards from all treatment groups are included (P < 0.001), but (B) not when individuals removed from the study due to symptomatic lead poisoning (all 100 mg/kg and 500 mg/kg treatment animals) are excluded (P = 0.432). Neither (C) sprint speed nor (D) balance were associated with final individual blood lead level (both P > 0.05; only animals in the Control, 1 mg/kg, and 10 mg/kg groups were measured).

**Tables**

**Table S1.** Summary of multiple linear regression models for factors associated with blood and bone lead levels of lizards from Sites B and F. Significant predictors at the α = 0.05 level are bolded.

|  | Predictor | Estimate | Std. error | t-value | p-value |
| --- | --- | --- | --- | --- | --- |
| Blood | Site | **-1083.90** | **179.80** | **-6.03** | **2.29e-8** |
|  | Sex | -279.50 | 169.80 | -1.65 | 0.10 |
|  | Site*sex | 271.00 | 231.20 | 1.17 | 0.24 |
| Bone | Site | **-33.06** | **12.57** | **-2.63** | **0.02** |
|  | SVL | 2.14 | 2.25 | 0.95 | 0.36 |
|  | Sex | -32.43 | 26.86 | -1.21 | 0.24 |

**Table S2.** Summary of negative binomial models for factors associated with balance in field collected animals. Significant predictors at the α = 0.05 level are bolded.

| Predictors | Estimate | Std. error | z-value | p-value |
| --- | --- | --- | --- | --- |
| Blood lead level | -2.02e-4 | 2.73e-6 | -1.09 | 0.28 |
| Sex | **-0.51** | **1.85e-4** | **-1.82** | **0.07** |
| Blood lead level*sex | 2.56e-4 | 3.02e-4 | 0.85 | 0.40 |
| Size | **-0.14** | **0.07** | **-2.13** | **0.03** |

**Table S3.** Summary of multiple regression model for factors associated with sprint speed in field collected animals. Significant predictors at the α = 0.05 level are bolded.

| Predictors | Estimate | Std. error | t-value | p-value |
| --- | --- | --- | --- | --- |
| Blood lead level | **1.12e-4** | **4.43e-5** | **2.54** | **0.01** |
| Sex | **2.50e-1** | **9.84e-2** | **2.54** | **0.01** |
| Blood lead level*sex | -1.89e-5 | 7.17e-5 | -0.26 | 0.80 |
| SVL | -1.20e-2 | 7.16e-3 | -1.67 | 0.09 |

**Table S4.** Summary of multiple linear regression models for factors associated with endurance in field collected animals. Significant predictors at the α = 0.05 level are bolded.

| Predictors | Estimate | Std. error | t-value | p-value |
| --- | --- | --- | --- | --- |
| Blood lead level | 0.04 | 0.02 | 1.83 | 0.09 |
| SVL | -1.82 | 1.91 | -0.95 | 0.35 |

**Table S5.** Mean blood lead level (µg/dL) with standard error (S.E.) of control and lead dosed treatment groups pre- and post-dosing, with post-dosing day of blood sample collection.

| Treatment group | Pre-dosing | Post-dosing | Post-dosing day |
| --- | --- | --- | --- |
| Control | 35.8 ± 7.25 | 78.6 ± 7.79 | Day 60 |
| 1 mg/kg | 34.8 ± 9.32 | 2232 ± 180 | Day 60 |
| 10 mg/kg | 46.5 ± 7.29 | 7684 ± 680 | Day 60 |
| 100 mg/kg | 17 ± 3.45 | 11500 ± 2616 | Day 30 |
| 500 mg/kg | 156 ± 128 | 21744 ± 7396 | Day 10 |

**Table S6.** Summary of two-way repeated measures ANOVA for predictors of lizard blood lead level. Significant predictors at the α = 0.05 level are bolded.

| Predictor | F-value | p-value |
| --- | --- | --- |
| Treatment group | **6.75** | **< 0.001** |
| Blood sample collection time | **34.32** | **< 0.001** |
| Treatment group*blood sample collection time | **6.54** | **< 0.001** |

**Table S7.** Summary of mixed effects model for factors associated with endurance among treatment groups during the 8-week period. Significant predictors at the α = 0.05 level are bolded.

|  | Predictor | Estimate | Std. error | t-value | p-value |
| --- | --- | --- | --- | --- | --- |
| Treatment group level | 1 mg/kg | 22.06 | 12.15 | 1.82 | 0.07 |
|  | 10 mg/kg | 7.66 | 12.15 | 0.63 | 0.53 |
|  | 100 mg/kg | **29.42** | **12.59** | **2.34** | **0.02** |
|  | 500 mg/kg | **51.52** | **14.53** | **3.54** | **< 0.01** |
|  | Run number | **9.70** | **3.34** | **2.09** | **< 0.01** |
|  | SVL | -1.20 | 0.94 | -1.28 | 0.20 |
|  | 1 mg/kg*run number | **-11.10** | **4.73** | **-2.35** | **0.02** |
|  | 10 mg/kg*run number | -3.93 | 4.73 | -0.83 | 0.41 |
|  | 100 mg/kg*run number | **-31.49** | **5.33** | **-5.91** | **< 0.001** |
|  | 500 mg/kg*run number | **-44.67** | **7.72** | **-5.79** | **< 0.001** |
| Individual level | Individual blood lead level | **-0.007** | **0.002** | **-3.40** | **< 0.01** |
|  | Run number | -0.09 | 2.89 | -0.03 | 0.97 |
|  | SVL | -0.98 | 1.01 | -0.98 | 0.34 |
|  | Blood*run number | **0.003** | **0.001** | **2.88** | **< 0.01** |

**Table S8.** Summary of multiple linear regression for factors associated with sprint speed following the 8-week dosing study.

|  | Predictor | Estimate | Std. error | t-value | p-value |
| --- | --- | --- | --- | --- | --- |
| Treatment group | 1 mg/kg | 0.17 | 0.11 | 1.41 | 0.17 |
|  | 10 mg /kg | 0.02 | 0.12 | 0.13 | 0.90 |
|  | SVL | -0.02 | 0.02 | -0.91 | 0.37 |
| Blood lead level | Blood lead | -3.76e-06 | 1.44e-05 | -0.26 | 0.80 |
|  | SVL | -1.50e-02 | 1.80e-02 | -0.84 | 0.41 |

**Table S9.** Summary of the general linear model for factors associated with lizard balance following the 8-week dosing study.

|  | Predictor | Estimate | Std. error | z-value | p-value |
| --- | --- | --- | --- | --- | --- |
| Treatment group | 1 mg/kg | 0.16 | 0.19 | 0.84 | 0.40 |
|  | 10 mg/kg | 0.17 | 0.20 | 0.85 | 0.40 |
|  | Size | -0.006 | 0.05 | -0.12 | 0.90 |
| Blood lead level | Blood lead | 6.38e-06 | 2.30e-05 | 0.28 | 0.78 |
|  | Size | -2.80e-02 | 4.82e-02 | -0.06 | 0.95 |

|  | **Term** | **P-value** | **Adjusted P-value** | **Odds Ratio** | **Combined Score** | **Genes** |
| --- | --- | --- | --- | --- | --- | --- |
|  | Fibroblast Growth Factor Receptor Binding (GO:0005104) | 0.003 | 0.140 | 29.71791 | 176.9859 | FGF19;FGF21 |
|  | Growth Factor Receptor Binding (GO:0070851) | 0.004 | 0.140 | 9.695992 | 52.46875 | PLSCR1;FGF19;FGF21 |
|  | O-acyltransferase Activity (GO:0008374) | 0.005 | 0.140 | 20.48585 | 108.0815 | SOAT2;PNPLA2 |
|  | Lipase Activity (GO:0016298) | 0.009 | 0.140 | 14.84403 | 69.53305 | ABHD2;PNPLA2 |
| **Molecular** | Phosphoric Diester Hydrolase Activity (GO:0008081) | 0.010 | 0.140 | 14.13575 | 64.94702 | PLCE1;GPCPD1 |
| **Function** | Cyclin-Dependent Protein Serine/Threonine Kinase Regulator Activity (GO:0016538) | 0.011 | 0.140 | 13.49186 | 60.83524 | CCNI2;CNPPD1 |
|  | Fructose Binding (GO:0070061) | 0.017 | 0.140 | 73.26103 | 297.9344 | PFKM |
|  | Retinyl-Palmitate Esterase Activity (GO:0050253) | 0.017 | 0.140 | 73.26103 | 297.9344 | PNPLA2 |
|  | Acetylcholine Receptor Inhibitor Activity (GO:0030550) | 0.017 | 0.140 | 73.26103 | 297.9344 | LY6E |
|  | Alcohol Dehydrogenase Activity, Zinc-Dependent (GO:0004024) | 0.021 | 0.140 | 58.60588 | 227.7499 | ADH7 |
|  | Intracellular Organelle Lumen (GO:0070013) | 0.009 | 0.219 | 2.951284 | 13.87653 | COL1A1;ALAS2;MUC1;CRISPLD2;  DGLUCY; OGN;HBA1;PNPLA2 |
|  | Apicolateral Plasma Membrane (GO:0016327) | 0.017 | 0.219 | 73.26103 | 297.9344 | MPDZ |
| **Cellular** | Serine/Threonine Protein Kinase Complex (GO:1902554) | 0.018 | 0.219 | 10.228 | 40.94643 | CCNI2;CNPPD1 |
| **Component** | Ficolin-1-Rich Granule Membrane (GO:0101003) | 0.018 | 0.219 | 10.228 | 40.94643 | PTPRN2;SLC11A1 |
|  | Ficolin-1-Rich Granule (GO:0101002) | 0.026 | 0.219 | 4.959819 | 18.14587 | PTPRN2;CRISPLD2;  SLC11A1 |
|  | Smc5-Smc6 Complex (GO:0030915) | 0.027 | 0.219 | 41.85714 | 150.7626 | NSMCE3 |
|  | Transcription Factor TFIIH Core Complex (GO:0000439) | 0.031 | 0.219 | 36.62316 | 127.6592 | GTF2H2 |
|  | Bicellular Tight Junction (GO:0005923) | 0.032 | 0.219 | 7.501228 | 25.84125 | MARVELD2;MPDZ |
|  | Transcription Factor TFIIH Holo Complex (GO:0005675) | 0.037 | 0.219 | 29.29559 | 96.33754 | GTF2H2 |
|  | Diacylglycerol Biosynthetic Process (GO:0006651) | 0.000 | 0.065 | 118.9612 | 989.7567 | PLCE1;PNPLA2 |
|  | Oxygen Transport (GO:0015671) | 0.001 | 0.065 | 74.33955 | 562.3452 | HBE1;HBA1 |
|  | Intestinal Lipid Absorption (GO:0098856) | 0.001 | 0.065 | 66.07629 | 486.7251 | FABP2;SOAT2 |
| **Biological** | Carbon Dioxide Transport (GO:0015670) | 0.001 | 0.065 | 66.07629 | 486.7251 | HBE1;HBA1 |
| **Process** | Intracellular Iron Ion Homeostasis (GO:0006879) | 0.001 | 0.065 | 19.23017 | 140.0898 | ALAS2;FRRS1;SLC11A1 |
|  | Gas Transport (GO:0015669) | 0.001 | 0.097 | 45.73594 | 307.7296 | HBE1;HBA1 |
|  | Glycerolipid Catabolic Process (GO:0046503) | 0.002 | 0.118 | 37.15485 | 236.253 | ABHD2;GPCPD1 |
|  | Glutamate Metabolic Process (GO:0006536) | 0.002 | 0.118 | 33.02322 | 202.9763 | TAT;DGLUCY |
|  | Hydrogen Peroxide Catabolic Process (GO:0042744) | 0.002 | 0.118 | 33.02322 | 202.9763 | HBE1;HBA1 |

**Table S10.** Gene ontology enrichment analysis of the differentially expressed genes in brain and liver tissues of brown anoles collected from a high lead versus a low lead contamination site. Genes that were upregulated in high lead lizards are indicated in red, genes that were downregulated are indicated in blue.

**Dataset S1 (separate file).**

Full master sheet of all blood lead data found during literature review. All studies had N ≥ 10 free-living adult individuals sampled.

**References S1**

1. C. E. Bryan, S. J. Christopher, B. C. Balmer, R. S. Wells, Establishing baseline levels of trace elements in blood and skin of bottlenose dolphins in Sarasota Bay, Florida: Implications for non-invasive monitoring. *Sci. Total Environ.* **388**, 325–342 (2007).
2. T. A. Rogers, “Lead exposure in large carnivores in the greater Yellowstone ecosystem,” The University of Montana. (2010).
3. V. Benito, *et al.,* Trace elements in blood collected from birds feeding in the area around Donana National Park affected by the toxic spill from the Aznalcollar mine. *Sci. Total Environ.* **242**, 309–323 (1999).
4. Dupont, *et al.,* Relationships between in vitro lymphoproliferative responses and levels of contaminants in blood of free-ranging adult harbour seals (*Phoca vitulina*) from the North Sea. *Aquat. Toxicol.* **142–143**, 210–220 (2013).
5. J. Burger, C. Jeitner, L. Schneider, R. Vogt, M. Gochfeld, Arsenic, cadmium, chromium, lead, mercury, and selenium levels in blood of four species of turtles from the Amazon in Brazil. *J. Toxicol. Environ. Heal. - Part A Curr. Issues.* **73**, 33–40 (2010).
6. Ráez-Bravo, *et al.,* Toxic and Essential Element Concentrations in Iberian Ibex (*Capra pyrenaica*) from the Sierra Nevada Natural Park (Spain): Reference intervals in whole blood. *Bull. Environ. Contam. Toxicol.* **96,** 273–280 (2016).
7. N. Ushine, *et al.,* Estimation of the reference lead (Pb) concentration levels affecting immune cells in the blood of black-headed gulls (Chroicocephalus ridibundus, Laridae). *Avian Conserv. Ecol.* **17** (2022).
8. M. Bucchia, *et al.,* Plasma levels of pollutants are much higher in loggerhead turtle populations from the Adriatic Sea than in those from open waters (Eastern Atlantic Ocean). *Sci. Total Environ.* **523,** 161–169 (2015).
9. G. Herring, C. A. Eagles-Smith, J. Goodell, J. A. Buck, J. J. Willacker, Small-mammal shooting as a conduit for lead exposure in avian scavengers. *Environ. Sci. Technol.* **55,** 12272–12280 (2021).
10. E. D. Sánchez, “Active and passive monitoring of lead poisoning in birds of prey in Spain,” University of Castilla-La Mancha. (2017).
11. M. Hernández, A. Margalida, Assessing the risk of lead exposure for the conservation of the endangered Pyrenean bearded vulture (*Gypaetus barbatus)* population. *Environ. Res.* **109,** 837–842 (2009).
12. J. Burger, *et al.,* Metal levels in blood, muscle and liver of water snakes (*Nerodia* spp.) from New Jersey, Tennessee and South Carolina. *Sci. Total Environ.* **373**, 556–563 (2007).
13. G. Herring, C. A. Eagles-Smith, D. E. Varland, Mercury and lead exposure in avian scavengers from the Pacific Northwest suggest risks to California condors: Implications for reintroduction and recovery. *Environ. Pollut.* **243,** 610–619 (2018).
14. Zavala-Félix, *et al.,* Trace elements concentration in blood of nesting Kemp’s Ridley turtles (*Lepidochelys kempii*) at Rancho Nuevo sanctuary, Tamaulipas, Mexico. *PLoS One* **17,** 1–12 (2022).
15. J. Rodriguez-Ramos, *et al.,* “Lead in griffon and cinereous vultures in central Spain: Correlations between clinical signs and blood lead levels” in *Ingestion of Lead from Spent Ammunition: Implications for Wildlife and Humans*, R. T. Watson, M. Fuller, M. Pokras, Eds. (The Peregrine Fund, 2009), pp. 235–236.
16. L. Gangoso, *et al.*, Blood lead levels in an endangered vulture decline following changes in hunting activity. *Environ. Res.* **252**, 0–2 (2024).
17. J. Burger, *et al.,* Arsenic, cadmium, chromium, lead, mercury and selenium concentrations in pine snakes (*Pituophis melanoleucus*) from the New Jersey pine barrens. *Arch. Environ. Contam. Toxicol.* **72,** 586–595 (2017).
18. L. G. Hivert, *et al.,* High blood lead concentrations in captive Tasmanian devils (*Sarcophilus harrisii*): a threat to the conservation of the species? *Aust. Vet. J.* **96,** 442–449 (2018).
19. M. Martinez-Haro, A. J. Green, R. Mateo, Effects of lead exposure on oxidative stress biomarkers and plasma biochemistry in waterbirds in the field. *Environ. Res.* **111**, 530–538 (2011).
20. E. Martínez-López, P. Gómez-Ramírez, S. Espín, M. P. Aldeguer, A. J. García-Fernández, Influence of a former mining area in the heavy metals concentrations in blood of free-living Mediterranean pond turtles (*Mauremys leprosa*). *Bull. Environ. Contam. Toxicol.* **99,** 167–172 (2017).
21. B. Fuchs, *et al.,* High concentrations of lead (Pb) in blood and milk of free-ranging brown bears (Ursus arctos) in Scandinavia. *Environ. Pollut.* **287** (2021).
22. S. C. McClelland, *et al.,* Sub-lethal exposure to lead is associated with heightened aggression in an urban songbird. *Sci. Total Environ.* **654,** 593–603 (2019).
23. K. Kucharska, Ł. J. Binkowski, G. Zaguła, K. Dudzik, Spatial, temporal and environmental differences in concentrations of lead in the blood of Mute swans from summer and winter sites in Poland. *Sci. Total Environ.* **830** (2022).
24. Sriram, W. Roe, M. Booth, B. Gartrell, Lead exposure in an urban, free-ranging parrot: Investigating prevalence, effect and source attribution using stable isotope analysis. *Sci. Total Environ.* **634,** 109–115 (2018).
25. E. Martínez-López, *et al.,* Blood δ-ALAD, lead and cadmium concentrations in spur-thighed tortoises (*Testudo graeca*) from southeastern Spain and northern Africa. *Ecotoxicology* **19,** 670–677 (2010).
26. T. Yazdanparast, *et al.,* Lead poisoning of backyard chickens: Implications for urban gardening and food production. *Environ. Pollut.* **310,** 119798 (2022).
27. R. Scheifler, *et al.*, Lead concentrations in feathers and blood of common blackbirds (*Turdus merula)* and in earthworms inhabiting unpolluted and moderately polluted urban areas. *Sci. Total Environ.* **371**, 197–205 (2006).
28. G. M. Wiemeyer, *et al.,* Repeated conservation threats across the Americas: High levels of blood and bone lead in the Andean condor widen the problem to a continental scale. *Environ. Pollut.* **220,** 672–679 (2017).
29. D. Rogival, J. Scheirs, W. De Coen, R. Verhagen, R. Blust, Metal blood levels and hematological characteristics in wood mice (*Apodemus sylvaticus L.*) along a metal pollution gradient. *Environ. Toxicol.* *Chem.* **25,** 149–157 (2006).
30. E. Guirlet, K. Das, M. Girondot, Maternal transfer of trace elements in leatherback turtles (*Dermochelys coriacea*) of French Guiana. *Aquat. Toxicol.* **88,** 267–276 (2008).
31. F. Ecke, *et al.,* Sublethal lead exposure alters movement behavior in free-ranging golden eagles. *Environ. Sci. Technol.* **51,** 5729–5736 (2017).
32. V. A. Slabe, *et al.*, Demographic implications of lead poisoning for eagles across North America. *Science.* **375** (2022).
33. M. Carneiro, *et al.,* Assessment of the exposure to heavy metals and arsenic in captive and free-living black kites (*Milvus migrans*) nesting in Portugal. *Ecotoxicol. Environ. Saf.* **160,** 191–196 (2018).
34. R. Brasso, *et al.,* Effects of lead exposure on birds breeding in the southeast Missouri lead mining district. USGS Sci. *Investig. Rep.* (2023).
35. V. Naidoo, K. Wolter, C. J. Botha, Lead ingestion as a potential contributing factor to the decline in vulture populations in southern Africa. *Environ. Res.* **152,** 150–156 (2017).
36. M. E. Church, *et al.,* Ammunition is the principal source of lead accumulated by California condors re-introduced to the wild. *Environ. Sci. Technol.* **40,** 6143–6150 (2006).
37. S. S. Lam, *et al.*, Lead concentrations in blood from incubating common eiders (Somateria mollissima) in the Baltic Sea. *Environ. Int.* **137** (2020).
38. J. L. Newth, *et al.,* Widespread exposure to lead affects the body condition of free-living whooper swans *Cygnus cygnus* wintering in Britain. *Environ. Pollut.* **209,** 60–67 (2016).
39. Z. P. Liu, Lead poisoning combined with cadmium in sheep and horses in the vicinity of non-ferrous metal smelters. *Sci. Total Environ.* **309,** 117–126 (2003).
40. L. van den Heever, H. Smit-Robinson, V. Naidoo, A. E. McKechnie, Blood and bone lead levels in South Africa’s Gyps vultures: Risk to nest-bound chicks and comparison with other avian taxa. *Sci. Total Environ.* **669,** 471–480 (2019).
41. M. M. Moore, *et al.*, Urban wall lizards are resilient to high levels of blood lead. *Environ. Res.* **264**, 120248 (2025).
42. M. M. Gillings, R. Ton, T. Harris, M. P. Taylor, S. C. Griffith, Blood lead increases and haemoglobin decreases in urban birds along a soil contamination gradient in a mining city. *Environ. Res.* **257**, 119236 (2024).
43. J. Mclelland, C. Reid, K. Mcinnes, W. Roe, Evidence of lead exposure in a free-ranging population of kea (*Nestor notabilis*). *Artic. J. Wildl. Dis.* **46,** 532–540 (2010).
44. J. Garcia-Fernandez, *et al.,* High levels of blood lead in griffon vultures (*Gyps fulvus*) from cazorla natural park (southern Spain). *Inc. Env. Toxicol* **20,** 459–463 (2005).
45. W. N. Beyer, *et al.,* Toxic exposure of songbirds to lead in the southeast Missouri lead mining district. *Arch. Environ. Contam. Toxicol.* **65,** 598–610 (2013).
46. F. M. Nilsen, *et al.,* Examining toxic trace element exposure in American alligators. *Environ. Int.* **128,** 324–334 (2019).
47. M. Adel, A. A. Cortés-gómez, M. Dadar, H. Riyahi, M. Girondot, A comparative study of inorganic elements in the blood of male and female Caspian pond turtles (*Mauremys caspica*) from the southern basin of the Caspian Sea. *Enrionmental Sci. Pollut. Res.* **24**, 24965–24979 (2017).
48. M. Humphries, J. Myburgh, R. Campbell, X. Combrink, High lead exposure and clinical signs of toxicosis in wild Nile crocodiles (*Crocodylus niloticus*) from a world heritage site: lake St Lucia estuarine system, South Africa. *Chemosphere* **303** (2022).
